# Integrity and miss-grouping as support for clusters in agglomerative hierarchical methods—the r-package octopucs

**DOI:** 10.1101/2024.08.01.606070

**Authors:** Ian MacGregor-Fors, Roger Guevara

## Abstract

The hierarchical clustering of communities based on species’compositional similarity (and abundance or frequency) is standard in community ecology to unveil large-scale patterns and underlay environmental causes of differentiation among communities. Often, the threshold to discretize clusters is arbitrary despite the existence of methods that minimize this bias. Most available techniques use the exact repeatability of clusters’ memberships under resampling protocols to define robust groups. Here, we propose a novel method to yield cluster support throughout the topology of hierarchical analyses. We acknowledge that the observed dataset may be biased. Instead of using the observed topology as a reference to work out the groups’ support, we compiled a consensus topology. Then, we borrowed the ecological concepts of reciprocal complementarities between a pair of communities and translated them into cluster integrity and contamination. This procedure allows for building support for groups even when there is a partial membership match after resampling the dataset. In addition, we present the R package octopucs in which we implemented the method reported here. Compared with other methods, the new proposal robustly detected changes in the group memberships, resulting in considerable differences in the pattern of supported clusters.

## INTRODUCTION

Identifying clusters of biotic communities with similar species composition and abundance is a significant goal in community ecology. Species composition and species abundance correlate with ecological processes (Cody and Diamond 1975, Morin 2011, Vellend 2016) and are sensitive to temporal and spatial heterogeneity (Proches 2005; Engelbrecht et al. 2007). Further, biotic communities respond to landscape and local habitat perturbations, local extinctions and invasions by exotic species (Stehn and Roland 2019), and global environmental changes (Bertrand et al. 2011). Systematic methods to estimate (dis)similarity among communities have received much attention from community ecologists. The early development of distance metrics between pairs of communities (e.g., Jaccard 1912) and key concepts such as beta-diversity (Whittaker 1960) followed the development of over 40 (dis)similarity indexes emphasizing different aspects of the spatial and temporal changes in species composition and their abundance among communities (Kolef et al. 2003). (Dis)Similarities matrixes are then processed through multivariate procedures and notoriously hierarchical agglomerative methods to uncover groups of similar communities. Contrary to (dis)similarity metrics and multivariate techniques, how to define the threshold at which groups are objectively identifiable has received less attention. Among some of the efforts made to provide objective methods to support emergent clusters in hierarchical agglomerative methods, a common target is the repeatability of the clusters’ memberships after resampling species with replacement (e.g., Hammer et al. 2001, Suzuki and Shimodaira 2006, Yu et al. 2018). Another method shuffles the observed memberships of joined clusters (keeping their sizes constant) to test for deviations from the random expectations in the similarity between the two groups (e.g., McKenna 2003). Yet another procedure seeks deviations in the correlation profiles between dissimilarities and their hierarchical order when species abundances are shuffled across sites at each division of the topology (e.g., Clarke et al. 2008).

Notwithstanding these efforts, we detected two drawbacks. These alternatives use the observed topology (and its subsets) as a reference, although biases may exist due to singularities in the database. Second, only perfect matches score positively in those methods aiming for cluster membership repeatability (e.g., Hammer et al. 2001, Suzuki and Shimodaira 2006, Yu et al. 2018). In these methods, there is no room for partial matches.

Here, we propose a novel method to define groups objectively in hierarchical agglomerative procedures. Similar to previous methods, our approach relies initially on resampling species. But instead of comparing topologies from every resampling event to the observed topology, we use them to put together the maximum likelihood hypothesis as a reference. Then, we weigh the compositional membership stability at every level of grouping. A given cluster may be regarded as unstable because it loses and gains members after resampling. Loosing and gaining members are, however, two distinct phenomena. In an analogy, a pile of apples is still a pile of apples, even when some are missing. In contrast, if some pears were mixed with the apples, the pile would no longer be a pile of apples, and the identity of the pile would change. The more pears appear in a pile, the more ambiguous their identity becomes.

Bearing that analogy in mind, at every resampling event, we collected information on losing and gaining members (against the most likely grouping hypothesis) at every level of clustering. We combined this information to build a support index. We compared the performance of this novel approach with one of the most widely used procedures, simprof (Clarke et al. 2008), to highlight the significant differences that will affect the conclusion of research relying on hierarchical agglomerative methods. Then, to facilitate the use of the proposed methodology, we implemented the R package octopucs, available at CRAN, and includes functions to implement our proposed methodology step by step and a conveniently nutshell-packed function, also named octopucs, for straightforward use. This implementation can use over 40 dis(similarity) metrics most used in community ecology.

## METHODS

### Background

After reviewing the currently available procedures to provide statistical support to clusters in HAC analyses, we are convinced that the challenge has yet to be addressed comprehensively, as the sole initial cluster topology is taken to be accurate and unbiased. Although some previous efforts considered the compositional membership of the clusters, they are based on a binary approach (exact match/no match) that weighs losing cluster members equally as gaining “no-members”. Such examples include Hammer et al. (2001) and Suzuki and Shimodaira (2006), who used bootstrap procedures on the whole dataset and multilevel bootstrap to obtain the probabilities of recovering identical groups.

The method proposed by Yu et al. (2018) seeks to establish a cutoff point in the topology. They used k-means cluster membership stability following resampling to define the number of supported clusters. Moreover, some approximations ignore the compositional membership of clusters but seek to identify departures from null model expectations. In the “similar profile approach” (simprof), the correlation profile observed between the matrix of similarities and grouping ranks is contrasted with the profile (95% confidence interval) obtained after permuting the dataset. Thus, the rank of grouping where the observed profile is higher than the random profile defines the cutoff point in the topology (cf. Clarke et al. 2008). Finally, McKenna (2003) used a Monte Carlo approach, randomizing the membership of connected clusters while keeping the group size constant to assess if each group’s observed (di)similarities differed from random expectations. A significant drawback of all these approximations is their reliability on output (the observed clustering hypothesis), which may reflect biases and uniqueness of the dataset.

### The rationale behind the proposed method

As noted, the output of HAC may be biased by the uniqueness of the dataset. We propose that if truly cohesive groups of sites are to be revealed by HAC, then their compositional memberships must suffer only minor variations when some of the species in the dataset (e.g., 20%) are randomly dropped (jackknifed), and a new HAC analysis is performed. Thus, by repeating this process many times, the outcomes can be put together in a consensus (or averaged) cluster hypothesis, which should minimize the potential artifacts of the original dataset. This is achieved by averaging the cophenetic distances (i.e., the branch length distance between any given pair of sites up to the nearest common node; Sneath and Sokal 1973) between every pair of sites over the whole set of the jackknife hypotheses. In addition, the procedure keeps track of the compositional membership at every level of the topology in each jackknife clustering hypothesis. This information is used to obtain the support metrics.

As part of the proposed procedure, we considered the following. For any level of grouping in the consensus hypothesis, there are only two theoretical outcomes: (a) the group is robust and genuinely exists (GE), that is, the membership is tight, or (b) there is a lack of cohesion in the membership of the group, and it does not exist as a robust group (1-GE). This consideration is not based on a given group’s size or compositional membership; instead, it is a logical *a priori* statement with a probability of 0.5 for either possibility. Subsequently, we contrasted the compositional membership for every group in the consensus hypothesis with the corresponding groups in each jackknife clustering hypothesis. Here, four outcomes are possible: (i) the exact group is recovered (nor gain or losses in the membership); (ii) all members of a given group in one of the jackknife clustering hypotheses are also clustered together in the corresponding group of the consensus hypothesis, but additional sites are intermingled in this group (gain of no member sites, contamination); (iii) a subset of the membership of a group in the consensus hypothesis is recovered in the jackknife clustering hypotheses (losing members or integrity); or (iv) as the previous one but mixed with additional sites (contamination and losing integrity). Hence, we defined two concepts to characterize each group in the consensus hypothesis over the whole set of jackknife clustering hypotheses: integrity (*I*), representing the probability of observing a group in the general hypothesis (*GH*) given that this group does exist *P*(*GH*|*GE*); and contamination (*C*), which is the probability of observing a group in the general hypothesis given that the group does not exist *P*(*GH*|-*GE*). The estimates of *I* and *C* were based on the complementarities in species composition between two communities (see Aguirre et al. 2010 for further details). According to the notation in Aguirre et al. (2010), *C*_*ab*_ and *C*_*ba*_ are the complementarities of site A to site B and vice versa, respectively. *I* is calculated as 1 – *C*_*ab*_, where the *C*_*ab*_ is the compositional complementary of a given cluster in the jackknife clustering hypotheses to the corresponding group in the consensus hypothesis. Similarly, the contamination, *C*, equals *C*_*ba*_, which is the compositional complementary of a cluster in the consensus clustering hypothesis to the corresponding group in the jackknife clustering hypotheses (Fig. 1). We calculated the conditional probability supporting each cluster as:

**Figure 1.**
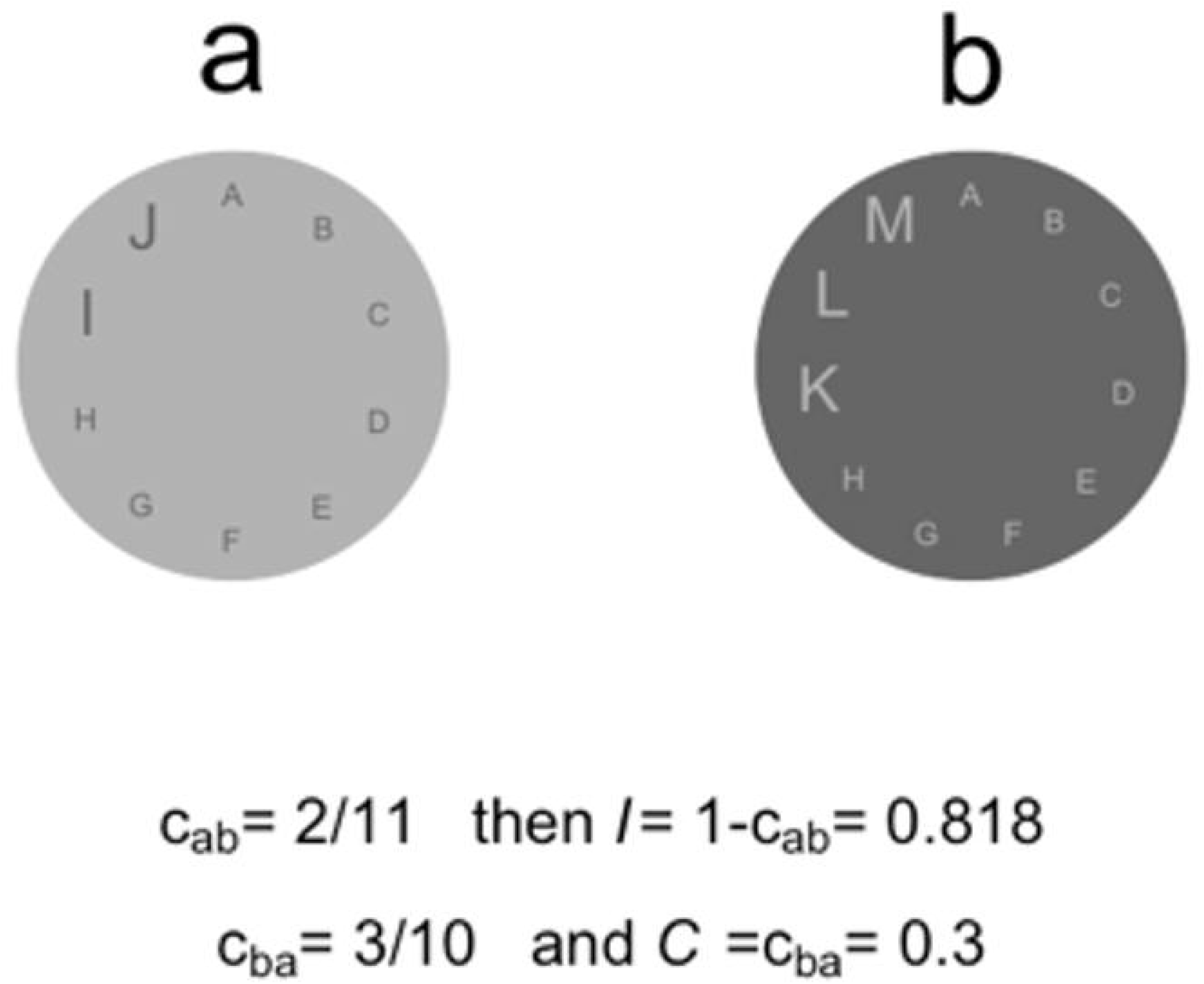
Schematic calculation of integrity (*I*) and contamination (*C*) of clusters in the general hypothesis (the average over several dendrograms obtained when 20% of the species are randomly removed). Assume cluster **a** is in one of the Jackknife dendrograms and cluster **b** is in the general hypothesis dendrogram. Then, the integrity of b relative to a is 0.818, as a contains two members not represented in **b**. Similarly, **b** has a 0.3 of contamination as it includes three members not present in **a**. *C*_*ab*_ and *C*_*ba*_ are the compositional complementarities between a given pair of groups.

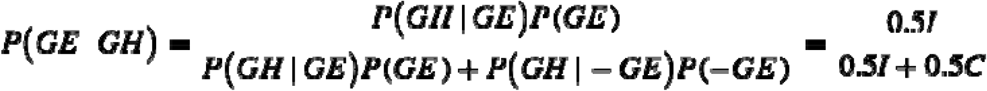

This procedure allowed for estimating the support for every cluster in the consensus hypothesis, regardless of its depth in the topology (i.e., supported groups would appear at different depths and with various internal configurations). For instance, a supported cluster that appears relatively high in the hierarchy may include equally supported nested subgroups, while some other subgroups may not be supported. The main difference between the proposed cluster Bayesian support and the procedures available currently (which are based on resampling of the data) is that rather than counting the frequency of the occurrence of a group with a rigorously defined membership (see Felsenstein 1985, Hammer et al. 2001, Suzuki and Shimodaira 2006, Yu et al. 2018), we weight two components, the cohesivity of a set of sites (defined here as integrity (I)) and the contamination (C) (the probability with which a cohesive set of sites is clustered together with other sites).

As a hypothetical example, when the same aggregation is recovered in all jackknife clustering hypotheses, integrity will equal one and C zero. When a subset of a given focal group of the consensus hypothesis is retrieved in the jackknife clustering hypotheses, I would be <1 and decrease with the subset’s size. However, if no members of other groups are incorporated, C will still equal zero. As members of different groups recover as a part of a focal group, the metric of contamination will be proportionally greater than zero, and 0.5 is the upper theoretical limit. To our knowledge, using these two components to support groups in HAC analyses is an innovation that may help ecologists (and scientists in other disciplines) interpret the outcome of HAC more objectively.

### Comparison with previous methods

We compared the output of our proposed method with that of Suzuki and Shimodaira (2006) (pvclust) using the R library of the same name and the similar profile method (simprof) (Clarke et al. 2008) using the functions available at https://rdrr.io/cran/clustsig/man/simprof.html. pvclust and simprof are two of the most widely used methods for HAC in ecology. We generated three datasets (Tables S1 and S2) for that purpose. The first database was a thoroughly random uniform distribution dataset of 30 sites and 20 species. The second database included three randomly generated groups with low internal variability but marked differences among the groups, with 30 sites and 20 species. The third database included two groups of highly variable samples.

In addition, we used a large data set of soil microbial (bacterial and fungal) OUTs in urban vacant lots of Yokohama, Kanagawa, Japan (Maehara et al. 2023; Maehara et al. 2024), and the dune meadow vegetation data (Jongman et al. 1987), dune, included in the R package vegan.

### The R package octopucs

We developed and distributed the package octopucs in CRAN (cran.r-project.org) to perform cluster support in HAC. This includes detailed code examples.

## RESULTS

A large proportion of the accepted metrics of distance and beta-diversity indexes that are used widely in community ecology (cf. Koleff et al. 2003) and other disciplines were implemented in octopucs, which has the advantage over other methods (e.g., pvclust and bootcluster) of only including some of the most common metrics (e.g., Euclidian, Manhattan, and correlation). The octopucs package effectively estimates the support at all levels of clustering, from pairs (support ranging between zero and one) up to the whole set of sites, where the support invariably equals one. Although pvclust works with some of the most relevant ecological metrics (e.g., Bray– Curtis), and the resulting topology of clustering is very similar to that of octopucs, support metrics for all nodes are zero (AU, BT, and chi-square values) (Fig. S1). This result opposed the cluster support performed in octopucs, the simprof, and the result of pvclust itself when the correlation distance (default option) was used (Fig. S1). However, the topology shows some variations (Fig. S1). All these were exemplified using the dune meadow vegetation data. Further comparisons were only performed between octopucs and simprof.

For the random uniform distribution dataset, octopucs and simprof performed well identifying no significant clusters (Fig. S2). For the dataset with relatively low intra-cluster variability, the cluster support in octopucs and simprof revealed the same three main groups (Fig. S3). Still, the main difference between octopucs and simprof was detecting a higher hierarchy cluster in octopucs containing two equally significant clusters.

For the customized database with two highly variable groups, considerable differences were observed between octopucs and simprof. While simprof recognized significant clusters of very low membership (two or even one samples), octopucs did not. Further, the only significant clusters of large memberships recognized by simprof were the least variable section of each group (Figure S4), also identified by octopucs. Most importantly, octopucs detected the two major groups as significant groups despite their high heterogeneity.

Further, the extensive database of microbial (fungi and bacteria) OTUs of vacant lots in Japan exemplified many differences between octopucs and simprof. Many low hierarchical clusters were supported by simprof, but only a few such clusters were supported by octopucs, including some coincidences between the methods (Figure 2). In contrast, most high hierarchical clusters supported by octopucs did not show in simprof. Finally, computational performance was also an advantage on the octopucs side as this procedure took 2.4 minutes to process the large database of microbial OTUs based on 1000 iterations, while for the same number of iterations, simprof took 4.4 hours (over a 110 times fold) in a computer with Apple-M2 Pro processor and 16 GB of memory.

**Figure 2.**
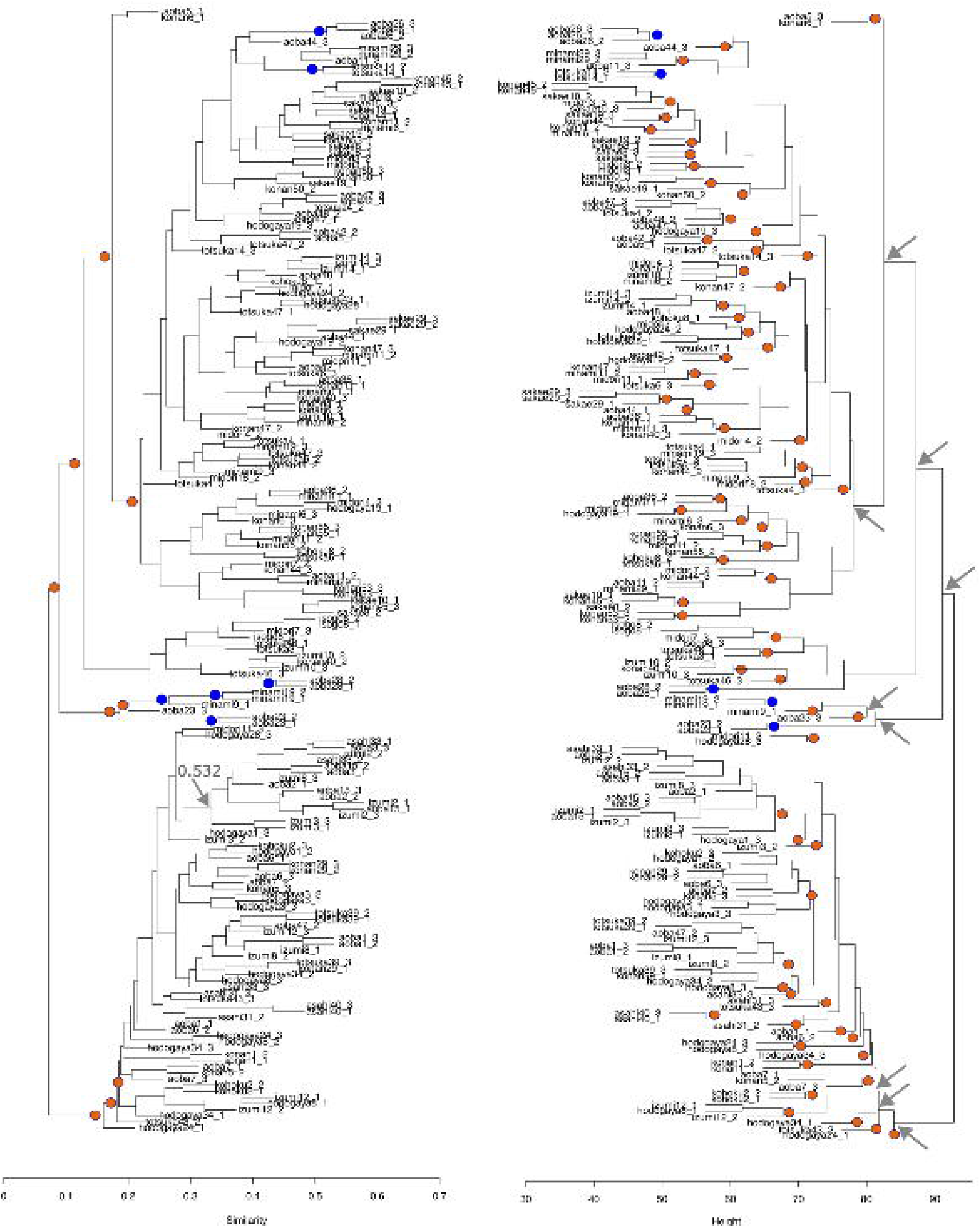
Comparative outputs produced by octopucs and the simprof method outputs for the OTUs of soil microorganisms (fungi and bacteria) in vacant lots in Yokohama, Kanagawa, Japan (Maehara et al. 2023; 2024). Red dots indicate clusters supported by one method, blue dots indicate clusters supported by both methods and arrows point to the main differences between the methods.

## DISCUSSION

Clustering communities based on the similarity of species composition and their abundances is widely used in community ecology. It is often the first step in the search for large-scale patterns and to infer the underlying causes of the differences among communities. Therefore, methods to discriminate among significant clusters must be robust and unbiased. Here, we showed that octopucs is a strong, reliable, and computationally efficient procedure that generates cluster support metrics in hierarchical agglomerative methods.

octopucs performed well in a variety of situations tested. This includes no detection of significant clusters in the customized random uniform dataset and detection of low and high variability clusters in other customized datasets. Further, octopucs showed limited support for low hierarchy clusters, which contrast markedly with a commonly used alternative, simprof, thus significantly reducing the potential overinterpretation of many small clusters. This fact highlights that not all clusters are meaningful, including some relatively populous ones. Under the octopucs algorithm, this can be interpreted as labile membership of clusters in the jackknifed clustering iterations. This issue appears exemplified in the analysis of microbial OTUs where a cluster of 15 observations is not supported by octopucs (support 0.52, integrity = 0.95; contamination =0.07), which contrasts with the support of the cluster by the simprof method. Instead, octopucs recognized this cluster as part of a large group (membership of almost 60 samples) that was not supported by simprof. This algorithm seems sensitive to the relatively low cohesion of groups since, in the example of microbial OTUs and in the customized dataset that included two noisy groups, the procedure fails to identify high hierarchy clusters recovered by octopucs.

Three sources of variation may explain the differences observed between the cluster Bayesian support method and the simprof method. The simprof method uses bootstrapping (resampling with replacement), whereas the proposed method uses jackknifing (sub-setting the database). In addition, simprof does not consider the cluster’s membership; instead, it relies on the correlation between the hierarchy of the clusters and the dissimilarity metric. The new proposal centers on the compositional membership (integrity and contamination) of the groups observed in a consensus clustering hypothesis compared with the corresponding clusters in each jackknife clustering hypothesis. The consensus clustering hypothesis is the third difference between the two methods.

The strength of detecting meaningful high hierarchic noisy clusters in octopucs is directly linked to the deep consideration that the algorithm does on membership identity at every level of clustering. Instead of a similarity estimate of samples in a cluster, octopucs quantifies the similarity between a group in the consensus grouping hypothesis and the corresponding group in each jackknifed cluster.

Here, we observed no effects of putting together a consensus grouping hypothesis that may minimize the uniqueness of the dataset. Topologies were rather similar or identical between octopucs and simprof. Nonetheless, this is a potential benefit of the method that could be shown in some specific cases.

In summary, octopucs offers a novel way to assess and interpret hierarchical agglomerative methods based on the combination of two metrics (i.e., integrity and contamination) according to the compositional memberships of the clusters. Because a wide array of distances is implemented in octopucs (including Euclidean, Manhattan, and Canberra), together with a large sample of beta-diversity indices, we foresee a significant contribution to the field of community ecology by helping to recognize objectively groups of sites based on species compositional similarities. Naturally, this procedure can be implemented in other areas of research where sites and their attributes can be arranged into an NxM matrix.

## Supporting information

https://drive.google.com/drive/folders/1I2h40xTuB0DlK0Uv7rnmj3xZ9magbyZU?usp=sharing

## ACKNOWLEDGMENTS

We thank Fabricio Villalobos for his valuable comments and suggestions, which enhanced the quality and clarity of the manuscript.

