## Supplementary material for "Integrity and miss-grouping as support for clusters in agglomerative hierarchical methods—the r-package octopucs": https://drive.google.com/drive/folders/1I2h40xTuB0DlK0Uv7rnmj3xZ9magbyZU?usp=sharing

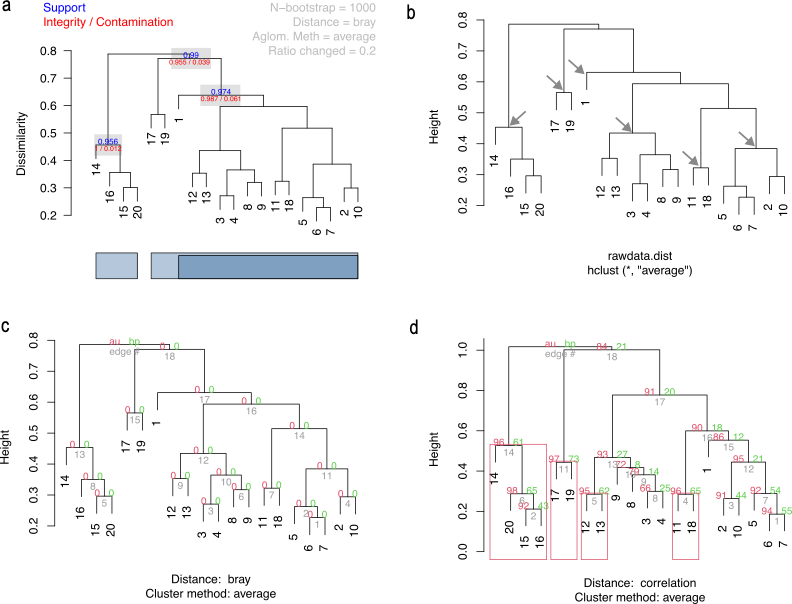

Figure S1. Comparison of clustering hypothesis produced by (a) octopucs, (b) simprof and pvclust with (c) Bray-Curtis, and (d) correlation distance. In the octopucs output, the figures (support, integrity, and contamination) indicate significant clusters. In the simprof result, we added arrows to indicate the significant clusters identified by the procedure. For the pvclust output, support metrics are approximately unbiased p values (au) and bootstrap probability values (BP); grey values are cluster identity.

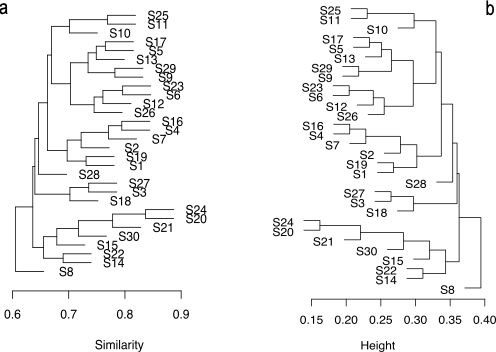

Figure S2. Comparison of clustering hypothesis on a customized databased following a random uniform distribution for which neither (a) octopucs nor (b) simprof detected significant clusters.

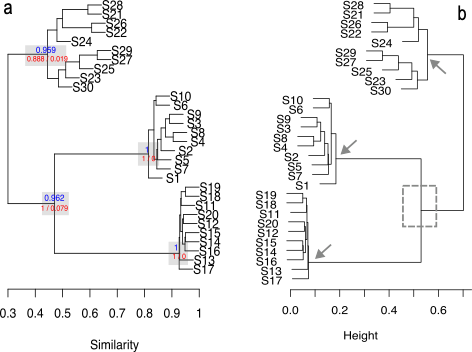

Figure S3. Comparison of clustering hypothesis on a customized databased produced by (a) octopucs, and (b) simprof highlighting the significant groups detected by each procedure, support metrics in octopucs and arrows in simprof, as well as the main difference in a high hierarchical cluster supported in octopucs but not in simprof (dashed rectangle).

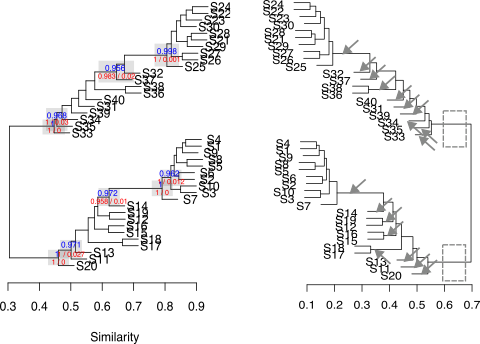

Figure S4. Comparison of clustering hypothesis on a customized databased produced by (a) octopucs, and (b) simprof highlighting the significant groups detected by each procedure, support metrics in octopucs and arrows in simprof, as well as the main differences in high hierarchical clusters supported in octopucs but not in simprof (dashed rectangles).
